# RNase 4 improves bottom-up modification mapping of *E. coli* total tRNAs using HILIC-MS/MS

**DOI:** 10.64898/2026.05.23.727428

**Authors:** Kaley M. Simcox, Maddy Zamecnik, Robert T. Kennedy, Kristin S. Koutmou

**Affiliations:** Department of Chemistry, University of Michigan, Ann Arbor, Michigan 48109, USA

**Keywords:** *Thraustochytrium* sp., orange pulp residue, omega-3 fatty acids, microalgae, circular economy

## Abstract

The structural and functional diversity of RNAs is expanded by the post-transcriptional incorporation of nucleoside variants. Emblematic of this, tRNAs contain extensive modifications that ensure their function during protein synthesis. Mass spectrometry has long been the field standard for identifying specific sites of chemical modifications on RNA. Nonetheless, mass spectrometry-based mapping approaches are not widely implemented. This is partially due to technical challenges associated with current methodologies including the limited diversity of available RNases, complexity of RNA mixtures, and conventional use ion-pairing reagents that require dedicated instrumentation. Here, we present a bottom-up liquid chromatography-tandem mass spectrometry (LC-MS/MS) workflow employing hydrophilic interaction liquid chromatography (HILIC) without ion-pairing reagents to globally map *E. coli* tRNA modifications. We implement orthogonal digestions using RNase 4 and a folded digestion scheme with RNase T1 to generate uniquely mappable oligonucleotides compatible with HILIC-MS/MS analysis and achieve 75-100% sequence coverage for most tRNA isoacceptors. HILIC-MS/MS matches the performance of traditional ion-pairing reverse-phased LC-MS/MS. This level of coverage allowed us to discover a new site of methylation (Gm17) in tRNA^Gly^, and confirm the presence of an s^4^U8 modification predicted in tRNA^Arg^. Furthermore, by applying this method to *E. coli* lacking the m^5^U54 methyltransferase (trmA) we confirmed the established dependence of acp^3^U47 insertion on m^5^U54 in tRNA^Phe^. Our findings show that RNase 4 improves bottom-up tRNA sequencing, enabling high-quality *E. coli* tRNA analysis without ion-pairing reagents.

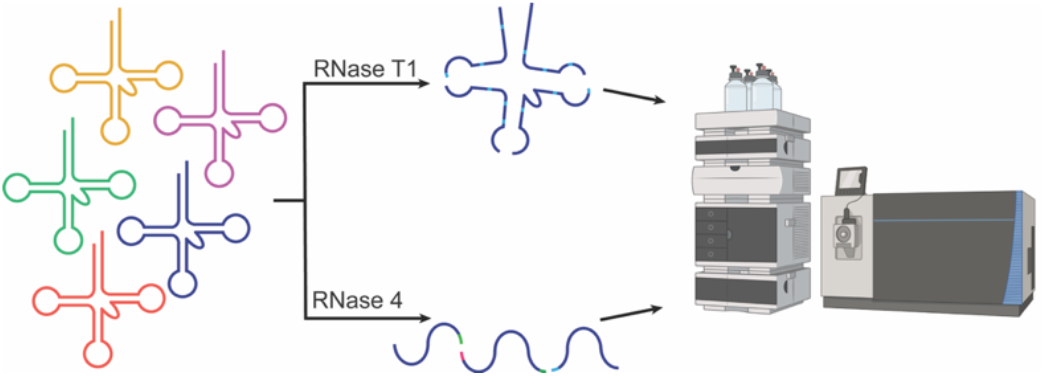

## INTRODUCTION

Across nature over 170 post-transcriptional chemical modifications are enzymatically installed into RNAs^1^. Transfer RNAs (tRNAs) are the most heavily modified class of RNAs, with an average of 10-20% of all tRNA nucleotides containing a modification^2,3^. These modifications influence the function of tRNAs by modulating their structure, stability and interactions on the ribosome^4–7^. While the importance of modifications for proper RNA function is universal, the identity and location of modifications vary between organisms^1^. Despite their clear importance and over half a century of study, the modification profiles of tRNAs remain to be fully determined for organisms beyond *E. coli* and *S. cerevisiae*^1^. This is in part due to the technical difficulties inherent in established liquid chromatography coupled to tandem mass spectrometry (LC-MS/MS) modification detection and mapping methods^8^. Nanopore direct RNA sequencing techniques are emerging as a powerful modern tool to predict the location of modifications in complex samples of RNA^9–12^. However, though these methods can predict where a modification exists, they are currently unable to directly determine the identity of a modification. As such, the further development of orthogonal methods capable of establishing the identity of modifications, such as LC-MS/MS, remains essential for the creation of robust RNA modification mapping pipelines.

Oligonucleotide LC-MS/MS enables the tandem identification and mapping of chemical modifications. In this approach, oligonucleotides are separated generally by length using liquid chromatography, then subjected to multiple MS-scans. The mass to charge ratio (m/z) of each oligonucleotide is first detected via MS1 scans, from which the intact mass can be calculated. Subsequently, each ion of interest undergoes a second MS/MS (or MS2) scan in which oligonucleotides are fragmented in the instrument, thereby enabling the positional determination of discrete chemical modifications based on the resulting fragment ions^13^. RNA fragmentation using collision-induced dissociation (CID) results in characteristic fragment ions based on the location on the phosphate backbone where the RNA was cleaved. In general, CID of RNAs have abundant C- and Y-type ions first characterized by Mcluckey in the 1990s^14^. Similarly to the field of proteomics, RNA modification mapping can be done using a ‘top-down’ approach where the full-length RNA is ionized into the MS and the fragment ions are used to map sites of modifications^15^. While top-down approaches have had success for oligonucleotide modification mapping of individual tRNAs, the low ionization efficiency of long RNA oligonucleotides limits this application to one tRNA species at a time^14,16^. For complex mixtures, such as total tRNA pools, ‘bottom-up’ approaches, where the RNA is enzymatically degraded to smaller lengths compatible with mass spectrometry and the unique oligonucleotides can be mapped back to a specific tRNA, are used^16–21^. Similar workflows are frequently employed in the field of proteomics^22,23^, but applying bottom-up modification mapping to RNA has been challenging due to limitations in the LC-MS/MS workflow and the lack of enzymes akin to proteases used for protein mapping. For RNA digestions prior to LC-MS/MS analysis, RNase T1 (guanosine (G) – specific) and RNase A (pyrimidine-specific) are the most used RNases. However, given that oligonucleotides contain many Gs and pyrimidines, these enzymes produce a myriad of short digestion products that are not unique to a specific tRNA, thus resulting in low sequence coverages^16,20,24^. This situation contrasts with the trypsin and non-trypsin (i.e. Asp-N, Glu-C) proteases used for protein bottom-up sequencing, which cleave a limited number of locations along a peptide backbone, allowing for the generation of peptides sufficiently long enough to be uniquely mapped^22,23^.

To improve RNA sequence coverage, previous work has focused on incorporating additional enzymes into the oligonucleotide digestion workflow and manipulating digestion schemes^18,19,24–27^. The use of multiple RNase enzymes capable of cleaving after different residues has been a particularly successful strategy to increase sequence coverage because these enzymes generate orthogonal digestion products for sequencing^8,16,20,24,25^. Combining the detected oligonucleotides digested by each enzyme enhances the sequence coverage and sequencing depth are improved across all RNAs. Some enzymes used to improve these workflows have a di- or tri-nucleotide consensus sequence that produce longer, more unique oligonucleotides that can be mapped back to specific sites on an RNA within a mixture^25–27^. Similarly, sequence coverages have also been improved using enzymes with shorter consensus sequences (e.g. U-specific and C-specific RNases)^20^. Further work has been done by manipulating the digestion schemes using RNase T1 and RNase A^16,27^. One study performed partial RNase T1 digestions which increased the length of RNA oligonucleotides, and improved sequence coverages across multiple mRNAs^27^. Another investigation found that using ‘folded’ tRNA RNase T1 and RNase A digestions improved sequence coverages for *S. cerevisiae* total tRNAs^16,25^. In this ‘folded’ digestion scheme, the tRNA is digested in the presence of magnesium and with a high salt concentration forcing the enzymes to primarily cleave in the single stranded regions of the RNA. This increased the length of the digestion products; thus, improving sequence coverages in *S. cerevisiae* total tRNA by having less non-unique digestion products^11,16^.

In bottom-up RNA sequencing schemes, the products of RNase digestion are analyzed by LC-MS/MS. This is typically accomplished using ion pairing reversed-phase (IP-RP) chromatography where ion pairing reagents (e.g. hexafluoroisopropanol and triethylamine) neutralize the negative charge of the phosphate backbone and enable oligonucleotide retention using reversed-phase chromatography. While this method has demonstrated superior resolution to alternative modes of chromatography, ion-pairing reagents linger in LC systems and their use requires dedicated instrumentation^28^. Thus, additional modes of chromatography are necessary for oligonucleotides analysis in areas where the use of PFAS are restricted or banned, in addition to labs without dedicated instrumentation (such as research cores). One alternative to IP-RP is hydrophilic interaction liquid chromatography (HILIC) without the use of ion-pairing reagents. In HILIC, oligonucleotides are thought to be retained through hydrogen bonding or electrostatic interactions between the RNA and the stationary phase. Therefore, oligonucleotides elute generally by length^19,28^. While HILIC generally has better resolution for shorter oligonucleotides (≤ 30 nt), this approach has been used successfully to directly map modifications in *S. cerevisiae* using RNase T1 and RNase A with high sequence coverages^11,16,29^.

This work presents methodological innovations designed to address challenges in the LC-MS/MS sequencing of complex modified RNA mixtures. Our sequencing scheme has the advantages of using only commercially available RNases and avoiding IP reagents, making it tractable for RNA scientists to conduct without dedicated instrumentation. The combined addition of a folded tRNA digestion scheme and inclusion of the ribonuclease IV (RNase 4) improve tRNA sequence coverages so that most tRNA isoacceptors have 70-100% sequence coverage. RNase 4 cleaves 3’ of uridines that are followed by a purine nucleotide. Comparing results to prior work shows that HILIC produces comparable results to IP-RP, negating the need for dedicated instrumentation and showing the applicability of our method for HILIC-MS/MS RNA modification mapping^20^. Additionally, we demonstrated the utility of this method by discovering a new site of modification in tRNA^Gly^, and applying it study of total tRNA purified from cells lacking the m^5^U54 methyltransferase, TrmA. We find evidence that 5-methyluridine (m^5^U) enhances the insertion of 3-(3-amino-3-carboxypropyl) uridine (acp^3^U), consistent with previous reports that m^5^U is required for other modifications to be fully incorporated by their modifying enzymes^30–34^. Overall, our findings reveal that the incorporation of RNase 4 into HILIC-MS/MS modification mapping workflows improves sequence coverages in *E. coli* total tRNA and provides a method with the sensitivity to detect RNA modification crosstalk.

## MATERIAL AND METHODS

### Cell growth and tRNA purifications

Wildtype (BW25113) and *trmA*Δ (JW3937) *E. coli* K-12 (Horizon Discovery) were grown in LB media at 37 °C overnight to saturation shaking at 250 rpm. *trmA*Δ cells were grown in the presence of kanamycin. Cells were harvested by centrifugation and nucleic acids were extracted as previously described with minor changes^35^. The cell pellet was resuspended in 20 mM sodium acetate and 20 mM magnesium acetate pH 5.5. RNA was extracted using acid phenol-chloroform isoamyl alcohol. The aqueous layer containing nucleic acids was brought to 20% isopropanol to precipitate DNA. The DNA pellet was discarded and the ribonucleic acids in the supernatant were precipitated by increasing the isopropanol to 60% by volume. The RNA was pelleted, resuspended in 10 mL of 200 mM tris-acetate pH 8.0, and incubated at 37 °C while shaking for 2 hours to deacylated the tRNAs. Deacylated tRNAs were ethanol precipitated overnight at −20 °C using 1/10^th^ volume 3 M sodium acetate pH 5.2, 2 μL glycoblue, and at least 2 volumes of 100% ethanol. The RNA was pelleted and resuspended in water. Total tRNA was purified from total RNA using a resourceQ strong anion exchange column on a FPLC. Mobile phase A was 300 mM NaCl and 25 mM sodium acetate pH 5.3. Mobile phase B was 800 mM NaCl and 25 mM sodium acetate. FPLC fractions containing tRNAs were identified using urea-PAGE with TBE buffer. Fractions containing only tRNA were pooled, precipitated overnight at – 20 °C using 1/10^th^ volume 3M sodium acetate pH 5.3, 2 μL glycoblue, and 3 volumes of 100% ethanol. The tRNA was pelleted and resuspended in 2 mL water. RNA concentration was determined using a Nanodrop spectrophotometer.

### Total tRNA RNase digestions

#### RNase T1 folded digestions

The purified *E. coli* total tRNA was digested using RNase T1 (ThermoFisher) as previously described^16^. Briefly, 315 μg of total tRNA in Milli-Q water was heated to 95 °C for 10 minutes. The RNA solution was incubated for 10 minutes at 37 ^°^C after adding 30 μL of 500 mM MES pH 6.0. Then, 3 μL of 2 M magnesium chloride was added to the RNA for a final concentration of 20 mM MgCl_2_. The tRNA solution was incubated at 37 °C for an additional 30 minutes. For RNase T1 digestions, 200 mM KCl was added to the RNA and digested using 6 U/μg RNase T1. After 1 hour incubation at 25 °C, the RNase T1 was removed using acid-phenol chloroform extractions. The aqueous layer containing the nucleic acid was washed with chloroform. To the washed aqueous layer, 0.13 U/μg calf intestinal phosphatase (CIP) was added and incubated for 1 hour at 25 °C. After 1 hour, the enzyme was removed using phenol chloroform extractions as previously described. The aqueous layer was ethanol precipitated overnight at −20 °C and resuspended in 7 μL of 200 mM LC-MS grade ammonium acetate prior to analysis.

#### RNase 4 complete digestions

Human RNase 4 (NEB) was buffer exchanged into 220 mM ammonium acetate using a Cytiva 5 mL HiTrap desalting column on an FPLC. Fractions containing the enzyme were concentrated using a 3 kDa MWCO filter. The purified *E. coli* total tRNA (315 μg) was digested in 220 mM ammonium acetate using 20 U/μg hRNase 4 for 2 hours at 37 °C. RNase 4 was removed using phenol chloroform extractions as previously described. To the washed aqueous layer, 5 U/μg T4 PNK was added and digested for 1 hour at 37 °C. After 1 hour incubation, T4 PNK was removed using phenol chloroform extractions. The washed aqueous phase was ethanol precipitated overnight at −20 °C and resuspended in 7 μL 200 mM LC-MS grade ammonium acetate.

### Total tRNA LC-MS/MS analysis

The digested total tRNA oligonucleotides were separated using a Waters ACQUITY Premier BEH Amide VanGuard FIT column (2.1 × 100 mm, with 1.7 μm particles containing 130 Å pores) using the LC conditions as previously described^4^. Briefly, 225 μg of the tRNA digest was injected on column. Mobile phase A was 25 mM ammonium acetate with 2.5 μM medronic acid in water and mobile phase B was 25 mM ammonium acetate with 2.5 μM medronic acid in 80% acetonitrile. The flow rate was 0.250 mL/min, the column set to 55 °C, and the autosampler was set to 4 °C.

An Agilent 1290 Infinity II Bio HPLC was interfaced to a Lumos Fusion Orbitrap mass spectrometer with a heated electrospray ionization source (HESI). Oligonucleotides were detected in negative ion mode with a spray voltage of 2900 V, with sheath and aux gases of 35 and 10 (arbitrary units), respectively. The ion transfer tube and vaporizer temperature were set to 350 °C. MS1 data was collected from 300-2000 m/z with charge states of 2-8 and the orbitrap resolution of 60K. The RF lens was set to 50% with an automated gain control (AGC) target of 480,000 and a normalized AGC target of 120%. Data was collected using data dependent acquisition where a minimum ion intensity of 25,000 was required to trigger fragmentation. Ions were fragmented using CID in the ion trap with a fixed collision energy of 35%. MS2 data was collected from 200-2000 m/z with an AGC target of 100,000, a normalized AGC target of 200%, and detected in the orbitrap with 30K resolution. After fragmentation ions with the same m/z were excluded for 3 seconds.

### LC-MS/MS data analysis

All data was analyzed using BioPharma Finder 5.1. *E. coli* tRNA sequences were obtained using MODOMICS^1^. *E. coli* tRNA modifications were programmed as building block modifications. Additionally, the use of the variable modification feature was employed. Here, we included the loss or gain of known *E. coli* tRNA modifications and 3’ linear, 2’-3’ cyclic phosphorylation and 5’ phosphorylation. Any detected 5’ phosphates were filtered to only include those mapped to the first nucleotide of a tRNA. The list of variable modifications and masses are listed in **SI Table 1**. Isobaric modifications such as pseudouridine do not generate unique fragment ions in oligonucleotide LC-MS/MS. To map pseudouridine sites, we relied on the known sites in Modomics. For RNase T1 and RNase 4 digestions, all masses observed were used in component detection, oligonucleotide identification requires MS/MS data, and a mass accuracy of 10 ppm was used. For RNase T1, custom specificity to cleave after inosine was added. The enzyme specificity level used was strict with no terminal 3’ phosphate. To account for the missed cleavages produced in the folded digestions, up to 20 missed cleavage sites were allowed. For RNase 4 analysis using BioPharma Finder, a nonspecific RNase was selected with the following custom specificity U-, D-. The various 3’ ends were searched for, 3’ linear, cyclic, and 3’ OH. Oligonucleotides were filtered to remove any non-unique oligonucleotides and Na^+^ adducts, resulting in only oligonucleotides that can be mapped back to a singular tRNA from the input sequences. A mass accuracy range of 10 ppm was used, and the minimum confidence score was set to 85%. The ASR was ≥ 1 and oligonucleotides without MS/MS data were removed.

### Nanopore library preparation and data analysis

Total tRNA from *E. coli* K-12 (Keio collection parental strain BW25113) and MRE600 strains were prepared for sequencing and analyzed as previously described^36,37^. In summary, tRNA were ligated to custom, pre-annealed 3’ and 5’ adapters ordered from IDT using T4 RNA ligase 2 and purified using RNAClean XP beads. The resulting RNA was then ligated to Oxford Nanopore’s RLA adapter using T4 DNA ligase and sequenced using RNA004 flow cells according to the manufacturer’s instructions and using the default settings.

After sequencing, the POD5 files were basecalled using Dorado 0.8.2 and ONT’s super accuracy^2^ model. The basecalled reads were aligned using BWA MEM version 0.7.19 and options “-W 13 -k 6 -x ont2d”. To reduce noise from high confidence alignments of reads to two different tRNAs, only a subset of distinct isodecoders were selected for the reference. Specifically, a fasta file was created for both strains using trnascan-1.4 and for, for each isoacceptor pool, the isoacceptor with the highest score appearing in both strains was selected for alignment. This sam file was then filtered with samtools version 1.21 to only primary, mapped reads with a mapping quality over 1. Finally, the program marginCaller was used to calculate the probability of a miscall at each position^38^.

## RESULTS

### Digestion of folded total *E. coli* tRNAs by RNase T1 produces RNA fragments appropriately sized for LC-MS/MS sequencing

We previously demonstrated the depth of *S. cerevisiae* total tRNA sequencing was significantly improved when the RNAs were folded prior to digestion with RNase T1^16^. RNase T1 cleaves the backbone of single-stranded RNAs on the 3’ side of every guanosine nucleotide (**Figure 1A, top**). Folding RNAs prior to digestion reduces the access of RNase T1 to single-stranded RNA regions, and yielded longer, more unique digestion products for sequencing. Given this observation, we posited that modification mapping of *E. coli* tRNAs might also be enhanced by digesting folded total tRNA. To evaluate if folding *E. coli* tRNAs prior to RNase T1 digestion also generates digestion products of sufficient length for sequencing, we examined the products of folded total *E. coli* RNA digestions. Due to the differences between the tRNA modification profiles we have observed in varying *E. coli* strains and environmental conditions by Nanopore direct RNA sequencing (**SI Figure 1**), we choose to examine total tRNA pools obtained from two different sources – either purified from MRE600 *E. coli* cells in our laboratory or purchased tRNA purified from MRE600 *E. coli* cells (Roche #10109541001). tRNA pools can vary between cell lines and growth conditions^39–41^, so we sought to discern if the source of RNA samples should be taken into consideration when selecting appropriate nuclease treatment conditions.

**Figure 1.**
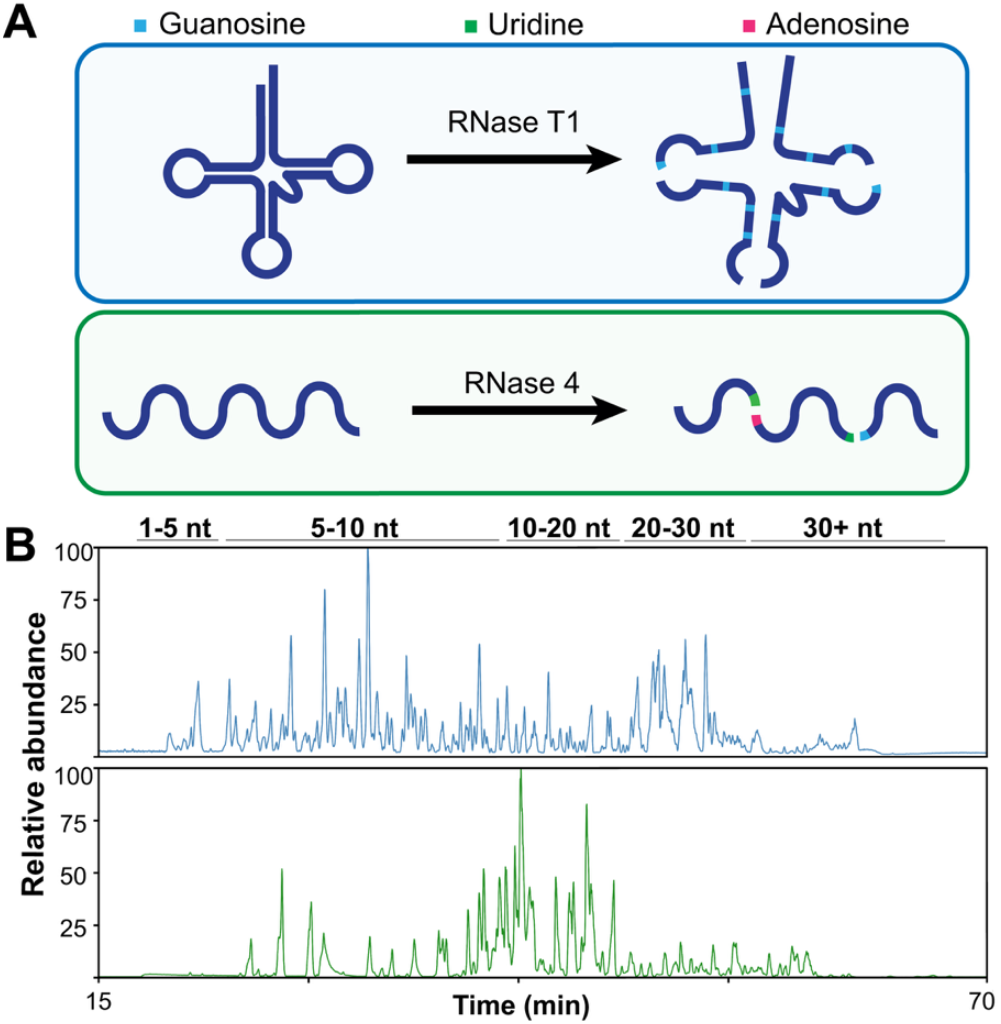
Total tRNA digestion schemes and HILIC-MS base peaks chromatograms of digested *E. coli* total tRNA. A) tRNA digestion schematic with RNase T1 (top) and RNase 4 (bottom). Folded RNase T1 digestions cleave after guanosines primarily in single-stranded regions of tRNAs. RNase 4 digestions cleave between U/A and U/G residues. B) HILIC-MS/MS base peak chromatograms of *E. coli* total tRNA digests using RNase T1 (blue trace) and RNase 4 (green trace). Relative abundance of MS signals was calculated by normalizing the signal to the maximum value in each trace.

We found notable differences in the digestion of the two samples. The purchased total tRNAs were digested to the same extent under a broad range of RNase T1 concentrations (25-6 U/ μg) (**SI Figure 2, right**). In contrast, the tRNA purified within our lab was much more sensitive to fluctuating concentrations of RNase T1. We determined the optimal amount of RNase T1 for purified *E. coli* total tRNA to be 6 U/μg. This concentration generates oligonucleotides with a broad range in lengths with a majority of digestion products in the range of 40-30 nt and ≤ 20 nt, whereas higher concentrations of RNase T1 produce oligonucleotides more closely reflecting complete digestion schemes with a majority of digestion products shorter than 10 nt (**SI Figure 2, left**). While the purchased tRNAs were less sensitive to increasing concentrations of RNase T1, the folded digestions scheme when applied to *E. coli* tRNAs resulted in oligonucleotides of comparable lengths to the method we developed for *S. cerevisiae* based on gel analysis of digestion products and LC-MS/MS analysis (**SI Figure 2**). We hypothesize that the differential sensitivities of in-house and commercially prepared tRNAs arise from increased salts in commercial tRNAs, which conceivably enhances tRNA folding resulting in fewer dynamic regions for RNase T1 to access^42^. The HILIC-UV chromatograms of *E. coli* total tRNA digest using 6 U/μg RNase T1 have a majority of the UV absorbance where oligonucleotides > 30 nt in length elute (**SI Figure 3, top**). However, the MS base peak chromatograms indicate detected oligonucleotides resulting from folded RNase T1 digestions are primarily between 5-10 nt or 20-30 nt (**Figure 1B, top panel; SI Figure 3, bottom**). The differences between UV and base peak chromatograms are most likely a result of the low ionization efficiency of longer oligonucleotides, in addition to ionization suppression resulting from co-elution of larger oligonucleotides. This is supported through the comparison of gel and LC-UV analysis as a majority of the signal corresponds to oligonucleotides with lengths ≥30 nt. (**SI Figures 2, 3**) These results suggest that the folded digestion scheme can be applied to any tRNA, or any highly structured RNA, by optimizing the amount of RNase.

### RNase 4 digestion of total *E. coli* tRNAs yields products with lengths compatible with LC-MS/MS

RNase 4 has been previously shown to improve modification mapping on a single mRNA transcript due to its dinucleotide consensus sequence^16^. However, the utility of RNase 4 for mapping modifications in a diverse mixture such as total tRNAs where the abundance and complexity of modifications are much larger has yet to be demonstrated. To examine if RNase 4 can lead to improved tRNA sequence coverages in LC-MS/MS workflows, we began by determining the concentration of RNase 4 necessary to digest *E. coli* tRNA into lengths compatible with LC-MS/MS. RNase 4 preferentially cleaves after uridines followed by purine nucleotides (A or G) (**Figure 1A, bottom**). We found that when digesting tRNAs with RNase 4 using the folded digestion scheme, many intact tRNAs were left (**SI Figure 4**). The presence of full-length tRNAs is problematic because it will result in poor HILIC resolution as current separations struggle to fully resolve longer oligonucleotides (>30 nt)^14,16,29^. This, coupled with the low ionization efficiency of longer oligonucleotides, would result in reduced MS signal for longer oligonucleotides and decreased sequence coverages. Thus, our data suggested the folded digestion scheme is not optimal for enzymes, like RNase 4, with dinucleotide or longer consensus sequence for tRNA modification mapping. However, when using an unfolded, or complete, digestion RNase 4 at 20 U/μg generated oligonucleotides compatible with LC-MS/MS workflows (**SI Figure 4**).

Base peak chromatograms of the complete RNase 4 digestion of *E. coli* total tRNA have a majority of the signal in the range where 10-20 mers elute (**Figure 1B, bottom**). However, the gel analysis and HILIC-UV chromatograms have a higher signal intensity for longer oligonucleotides (25-40 mers) (**SI Figure 5, top**). As seen in the RNase T1 results, these differences in signal intensity can be attributed to the decreased HILIC resolution of longer oligonucleotides and the decreased ionization efficiency of longer oligonucleotides. Both occurring together result in a higher base peak chromatogram signal in the range of 10-20 mers for RNase 4 digestion. Regardless, all three techniques are in agreement that RNase 4 digests *E. coli* total tRNA into lengths compatible with LC-MS/MS workflows.

### RNase T1 folded digestion results in long oligonucleotides for modification mapping

After obtaining *E. coli* total tRNA samples, we moved on to sequencing samples by HILIC-MS/MS. Oligonucleotides ranging from 2-42 nt were identified using BioPharma Finder from folded RNase T1 digestion of *E. coli* total tRNA (**SI Table 2-3**). Of these oligonucleotides, 21.7% were nonunique oligonucleotides that map back to positions on multiple tRNAs. These short, nonunique oligonucleotides ranged in length from 2-18 nt (**SI Table 2**). While a majority of the nonunique oligonucleotides are generated when RNase T1 cleaves at every possible G (no missed cleavages), a handful result when RNase T1 misses a cleavage opportunity. Of these, the oligonucleotides that are ≥ 4 nt in length exclusively map to positions within the D-loop or T-loop (**SI Table 2**). This is a result of the high sequence similarity within these tRNA regions. Therefore, to produce a unique oligonucleotide that maps to those regions, additional missed cleavages from RNase T1 are necessary to generate an oligonucleotide that does not map to multiple tRNAs. This reliance on missed cleavages limits the utility RNase T1, as for some tRNA isoacceptors families the generation of unique oligonucleotides would require the enzyme to miss > 6 potential cleavage sites. Due to the ambiguity of which tRNA many of the digested oligonucleotides originate from, the nonunique oligonucleotides that mapped to multiple tRNAs were excluded from all sequence coverage calculations. The remaining unique oligonucleotides varied in length from 4-42 nt, with the average length of 14 nt (**SI Table 3**). The distribution of the unique digestion product lengths are as follows: 20.1% ≤ 7 nt, 64.9% between 8-20 nt, and 15.0% ≥ 21 nt. Compared to a complete digestion of *E. coli* total tRNA, we would expect 164 of unique digestion products ranging from 3-20 nt with an average unique oligonucleotide having 7.5 residues. Thus, the folded digestions increase the length of digested oligonucleotides on average, consequently improving sequence coverages (**SI Table 3-4, 7**).

Our previous work with *S. cerevisiae* showed that when using the folded digestion scheme, we detected oligonucleotides with no missed cleavages and many with multiple missed cleavages^16^. This was also observed in the *E. coli* RNase T1 data. Exemplary of this, twenty oligonucleotides were detected that map back to tRNA^Ini^ with 85.7% sequence coverage (**Figure 2**). Of the detected oligonucleotides, eight had zero missed cleavages, one had three missed cleavages, two had four missed cleavages, and nine had 5+ missed cleavages (**Figure 2, SI Table 3-4**). These oligonucleotides with missed cleavages are on average much longer than those without missed, thus facilitating mapping to areas of the tRNA that are G-rich and improving sequence coverages.

**Figure 2.**
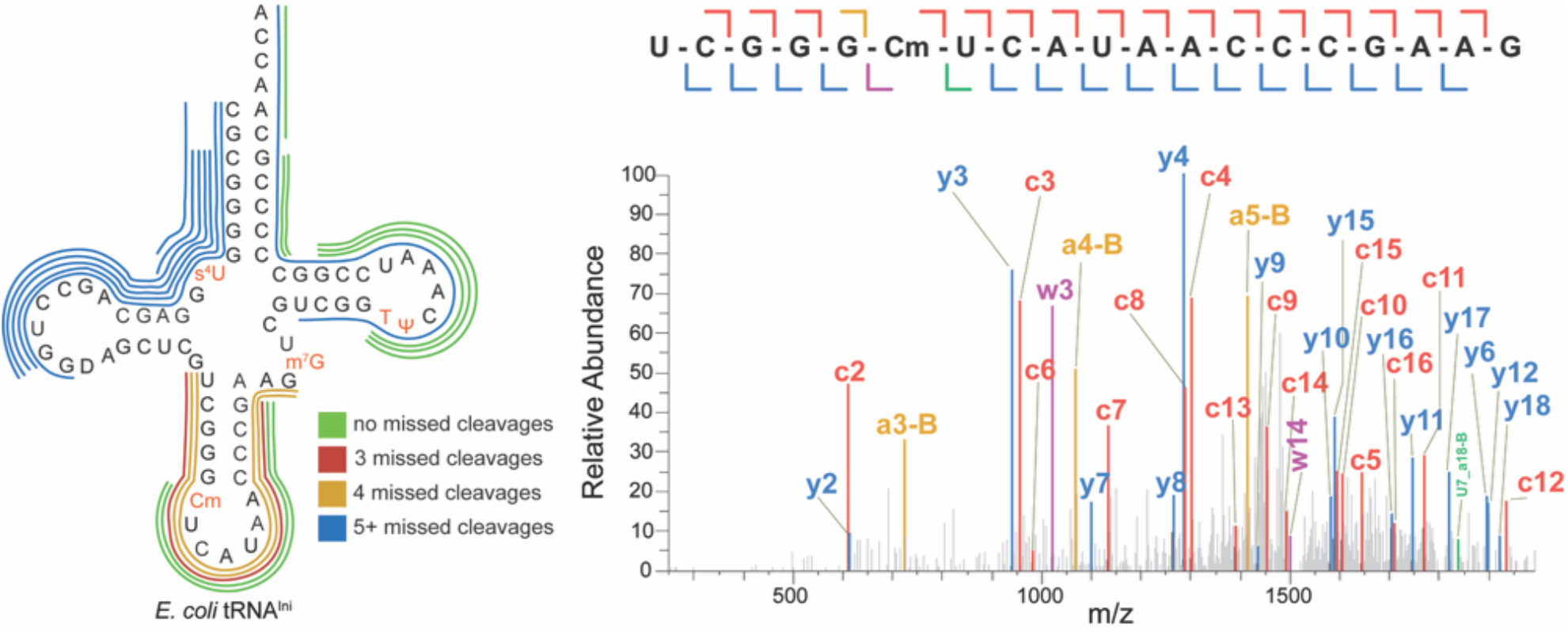
Sequence coverage of tRNA^Ini^ from RNase T1 digestion. Left - Detected tRN^Ini^ oligonucleotides in RNase T1 digestion. Each line represents a single oligonucleotide. The color of the line corresponds to the number of missed cleavages as a result of the folded digestion scheme. No missed cleavages (green), 3 missed cleavages (red), four missed cleavages (yellow), and five or more (blue). Right – MS/MS spectra of tRNA oligonucleotide with m/z 1517.21 that maps to the anticodon stemloop with 4 missed cleavages. The colors in the MS/MS spectra correspond to the type of fragment ion detected. C-type (red), a-B type (yellow), Y-type (blue), W-type (purple), internal fragment (green).

### RNase 4 generates unique oligonucleotides to enhance modification mapping

After digesting *E. coli* total tRNA with RNase 4, the resulting oligonucleotides were analyzed using LC-MS/MS. Of the oligonucleotides identified, less than 10% oligonucleotides were short, nonunique oligonucleotides that could not be mapped to a specific tRNA in either biological replicate. Like the RNase T1 digest, these shorter oligonucleotides primarily map back to sites of high sequence similarities (e.g. D-loop, acceptor stem) and were excluded from sequence coverage calculations. The oligonucleotides were primarily cleaved after a uridine or dihydrouridine followed by a purine (A or G) (**SI Table 5**). However, ∼20% of detected oligonucleotides were cleaved after U or D were followed by a pyrimidine nucleotide (U or C) (**Figure 3A**). These results support previous work showing that RNase 4 preferentially cleaves U/A and U/G residues, followed by a lower preference for U/C, or U/U residues and can tolerate chemical modifications of uridine (i.e. D)^26^.

**Figure 3.**
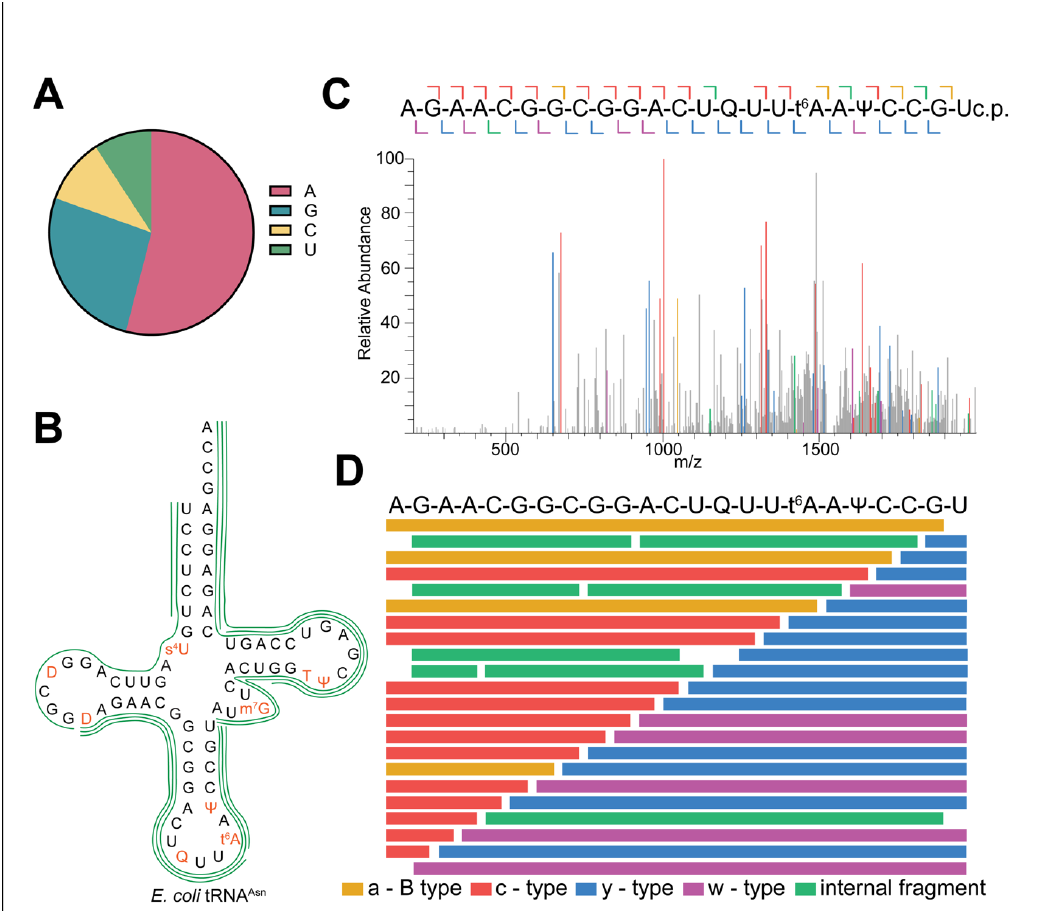
Sequence coverage of tRNA^Asn^ from RNase 4 digestion. A) Percentage of RNase 4 cleavage in detected oligonucleotides from *E. coli* total tRNA. A, G, C, and U in represent the 5’ terminal nucleotide resulting from RNase 4 digestion, excluding those that map to tRNA nucleotide position 1. B) RNase 4 sequence coverage of tRNA^Asn^. Each line corresponds to a detected oligonucleotide. C) MS/MS spectrum of tRNA^Asn^ oligonucleotide that maps from the D-loop across the anticodon stem loop. This oligonucleotide has a m/z of 1542.815 and a terminal 3’ cyclic phosphate. The colors of the fragment ions correspond to the ion type generated through ion-trap CID (shown in panel D). D) Fragmentation map of tRNA^Asn^ oligonucleotide from panel C.

Previous reports found that removing the terminal phosphate from RNAs prior to LC-MS/MS analysis improves spectral quality^43,44^. As such, the terminal phosphate is commonly removed using a phosphatase, such as bacterial alkaline phosphatase. However, RNase 4 produces a mix of 3’ linear phosphates and cyclic phosphates. Previously, T4 PNK was used to remove the heterogenous phosphate ends resulting from digestion with RNase 4^26^. In this study, we used T4 PNK to remove the 3’ phosphates; however, the oligonucleotides identified have a mix of 3’ ends containing linear phosphates, 2’-3’ cyclic phosphates, and 3’ OH ends (**SI Table 5**). This suggests that the amount of T4 PNK or length of T4 PNK digestion needs to be optimized in future experiments for tRNA mapping. Despite heterogeneity in the 3’ ends, fragmentation of these oligonucleotides resulted in sufficient abundance of sequence informative fragment ions. As such, oligonucleotides generated through RNase 4 digestion resulted in high sequence coverage across many *E. coli* tRNA isoacceptors (**Figure 3B-D, SI Table 5-7**). Emblematic of this, the coverage of tRNA^Asn^ was improved from 23.7% with RNase T1, to 100% with RNase 4 (**Figure 3B**). Furthermore, in general RNase 4 allowed oligonucleotides mapped to the anticodon loop captured labile modifications such as t^6^A and queuosine, which can fragment in the MS source and skew MS/MS data analysis^16,45^ (**Figure 3C-D**).

### Folded RNase digestion produced heterogeneous 3’ ends and 5’ ends

The variable modification feature in BioPharma Finder enables the identification of RNA modifications without prior knowledge of their position on an RNA. Here, the gain and loss of a modification on its target nucleobase used to identify new sites of modifications. **SI Table 1** lists the variable modifications used and the nucleobase they can modify. In addition to base modifications, using this feature led to the discovery of heterogenous 3’ ends in the folded RNase T1 digestion, where detected oligonucleotides had a shift in mass in the W and Y type ions corresponding to the mass of 2’ 3’-cyclic (+61.9 Da) and 3’ phosphates (+79.9 Da) (**SI Figure 6, SI Table 3**). While previously unreported in our folded digestion scheme, this is an unsurprising result as the folded digestions are a type of partial RNase T1 digest which have been known to generate a mix of phosphate ends^27^. As such, we have reanalyzed data from folded RNase T1 digestions of *S. cerevisiae* total tRNA and have identified variable 3’ ends (**SI Table 7**). This suggests that the variable phosphate ends are a result of the folded digestion scheme, rather than specific to the tRNA analyzed. Furthermore, incorporating the variable phosphate ends in our data analysis consequently improved coverages across many tRNAs by increasing the number of detected features, especially across regions of low coverage such as the D-loop (**SI Table 8**). Further work can be done to optimize the amount of phosphatase used or implement the use of T4 PNK which is able of cleaving cyclic phosphates to reduce the abundance of these oligonucleotides. However, similar to the RNase 4 digestions, even with the presence of 3’ phosphate ends on the digested oligonucleotides high abundant sequence informative fragment ions used are generated.

In addition to variable 3’ ends, we detected 5’ terminal phosphates on the 1^st^ nucleotide of a tRNA oligonucleotide in both E. coli total tRNA RNase digestions (**SI Table 3, 5**). C-type fragment ions are shifted corresponding to the mass of a phosphate (+79.9 Da) (SI Figure 7). During tRNA maturation, RNase P cleaves the 5’ leader sequence, resulting in a terminal 5’ phosphate on mature tRNAs^46^. Due to the addition of a phosphatase in our digestions, we would expect this 5’ terminal phosphate to be cleaved resulting in a 5’-OH. However, the presence of 5’ phosphates are likely a result of residual phenol-chloroform from the RNase extraction prior to phosphatase digestion. Here, if there is any residual phenol chloroform we would expect phosphatase activity to decrease resulting in a mixture of hydroxyl and phosphate groups to reside on the 5’ end of a nucleotide in position 1. The identification of these phosphorylated oligonucleotides increased the number of detected oligonucleotides increased RNase T1 and RNase 4 sequence coverages and depth compared to our previously reported analysis which does not include variable phosphate ends (**SI Table 3-6**).

### Variable modifications reveal new sites of tRNA modification

Using this adaptive workflow to map tRNA modifications, we recapitulated previous findings that cmo^5^U34 is further modified to cmo^5^Um34 when cells are grown to stationary phase on tRNAs Ala^VGC^ and Ser^UGA^ (**SI Figure 8 and Table 3**)^47^. This is evident as the fragment ions containing cmo5U34 have a shift in mass corresponding to the addition of a methyl group (+14 Da) (SI figure 8). Additionally, we detected oligonucleotides with and without modifications at known positions, indicating that these modifications are not fully modified on each copy of a tRNA. Some of these have been previously reported such as s^4^U8 on tRNA Glu^{UC^ and Gly^GCC 48^. The other modifications we detected to be partially modified are anticodon modifications such as t^6^A37 and mnm^5^U34. While these modifications are larger and can fragment in the source of a mass spectrometer, we are confident that these are a result of substiochiometeric modification at these sites due to differences in HILIC retention times (**SI Figure 9, SI Table 3, 5**). In addition to supporting previous findings about tRNA modification biology, we have also confirmed the presence of s^4^U8 on tRNA Arg^UCU^ predicted by RT-based assay, but previously not confirmed using mass spectrometry^48^ (**Figure 4**). Here the fragment ions containing U8 have a characteristic shift in mass of +15.9 Da, corresponding to the presence of s^4^U8 on tRNA Arg^UCU^ (**Figure 4A**). Furthermore, total tRNA from both a wild type cell line and a knockout cell line for *ThiI*, the enzyme that incorporates s^4^U8 in *E. coli* tRNAs, were sequenced using nanopore direct RNA sequencing. The tRNA from the knockout cell line produced a lower error rate at site 8 in tRNA Arg^UCU^, characteristic of a change in modification status at that position in the knockout (**Figure 4B**). This suggests that U8 in tRNA Arg^UCU^ is natively modified to s^4^U, and agrees with the previous RT-based assay prediction of s^4^U8 on tRNA Arg^UCU^ and the LC-MS/MS data collected here.

**Figure 4.**
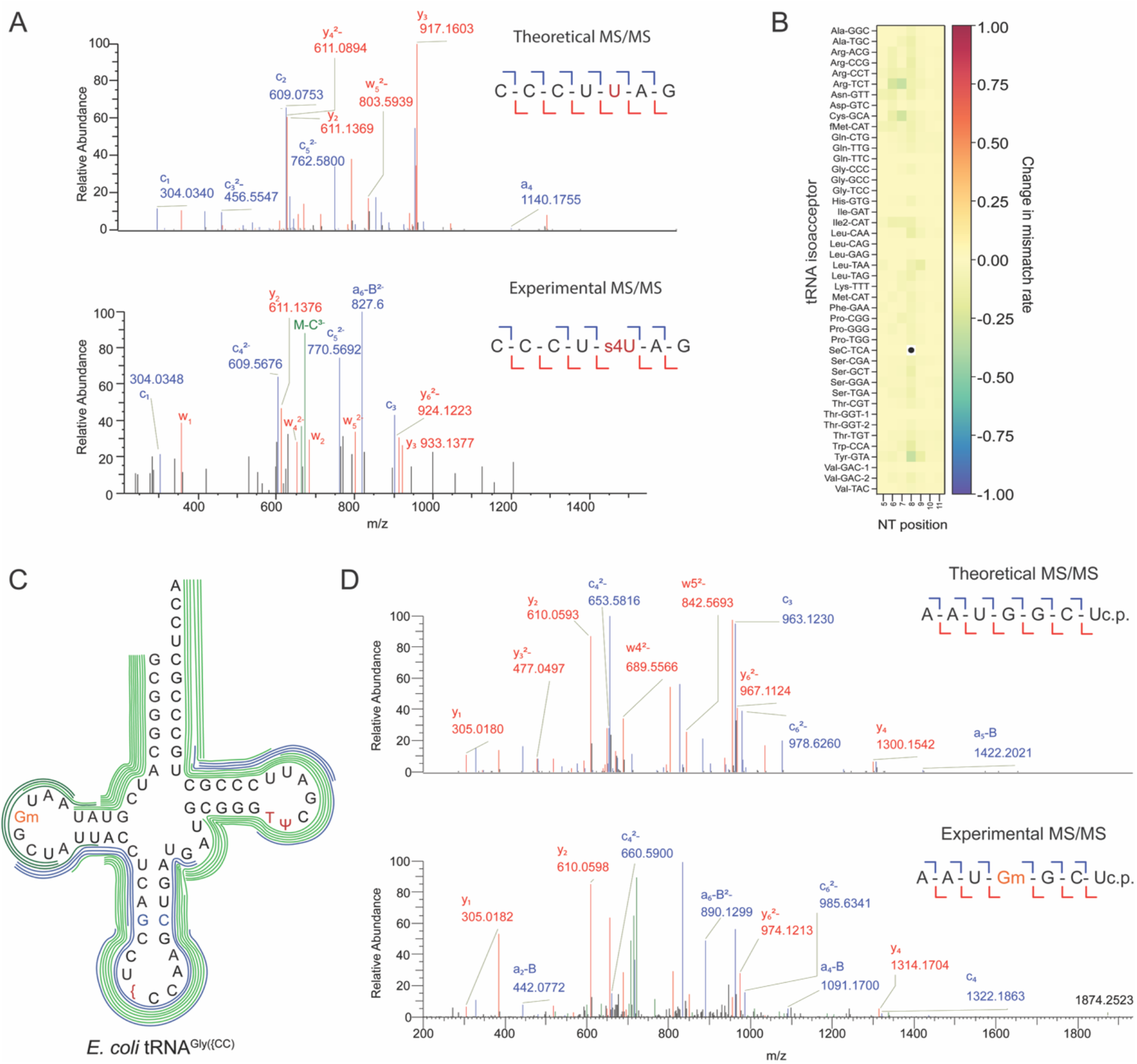
New sites of RNA modification on tRNA Arg^UCU^ and Gly^{CC^. A) MS/MS spectra from a tRNA oligonucleotide mapping s^4^U8 to tRNA Arg^UCU^. Top – theoretical MS/MS spectral of with U8. Bottom – experimental MS/MS spectrum from oligonucleotide with m/z 717.760 mapping s^4^U8 on tRNA Arg^UCU^. The shifts in mass correspond to the mass of the s^4^U modification (+15.9 Da) seen in the shifts in the Y^3^ and C^5^ ions. B) Nanopore direct RNA sequencing base-miscall difference from WT and thiI KO heatmap. C) Detected tRNA Gly^{CC^ ({ = 5-methylaminomethyluridine (mnm5U)) oligonucleotides from RNase T1 (blue) and RNase 4 (green) digestion. In Modomics, position 17 is noted as an unmodified guanosine^1^. All detected oligonucleotides in with coverage at G17 map to Gm17. This is observed in the trmA≥ tRNA digests in SI Table 11. D) Top – theoretical MS/MS spectra of oligonucleotide mapping G17. Bottom – experimental MS/MS spectra of oligonucleotide with m/z 758.763 mapping Gm17 to tRNA Gly^{CC^. The shift in mass corresponding to the methylation (+14 Da) is observed in the C_4_, C_6_ and Y_4_ and Y_6_ions.

Moreover, we detected a new site of tRNA modification on tRNA Gly^mnm5UCC^. Across many *E. coli* tRNA isoacceptors, G17 is known to be modified by trmH to 2’-O-methylguanosine (Gm). Our data shows that all detected oligonucleotides with coverage of G17 in tRNA Gly^k2CCC^ are modified to Gm17 (**Figure 4C-D**). Here, the fragment ions with G17 have a +14 Da shift in mass, representative of the addition of a methyl group. While we are unable to differentiate positional isomers in this method, many tRNAs are modified to Gm at position 17, which leads us to believe tRNA Gly^mnm5UCC^ contains Gm17 rather than different methylated guanosine. To our knowledge, Gly^mnm5UCCC^ is not annotated to have a modification at position 17; thus revealing a new site of modification for this tRNA. Together these results show the utility of our workflow to achieve high sequence coverage and uncover novel sites of tRNA modification.

### The combination of RNase T1 and RNase 4 detected oligonucleotides improves sequence coverages

The average individual sequence coverages for RNase T1 and RNase 4 across the *E. coli* tRNA isoacceptors were 65.8% and 57.1%, respectively (**SI Table 9**). Despite the similar average values, when the sequence coverages are combined, the inclusion of RNase 4 improves RNase T1 sequence coverages on average by ∼14%. Furthermore, the combination of folded RNase T1 digestion and RNase 4 resulted a sequence coverage ranging from 75-100% for a majority of tRNA isoacceptors (**Figure 5**).

**Figure 5.**
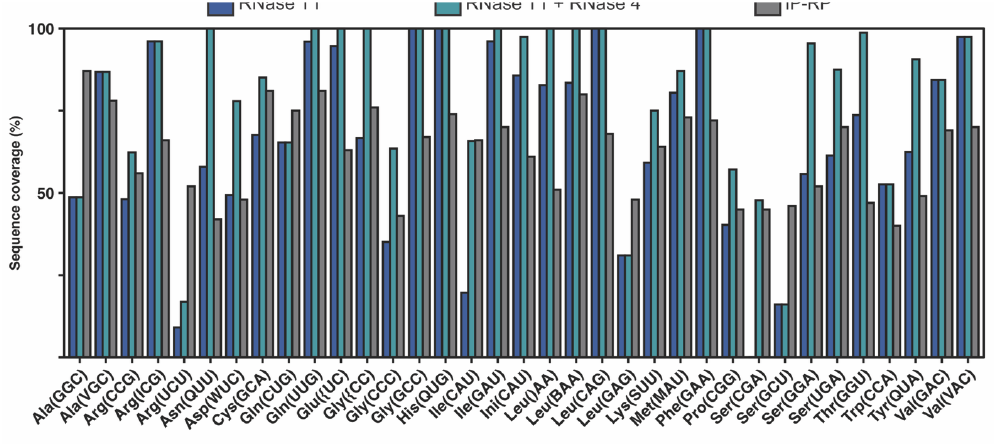
*E. coli* total tRNA sequence coverages. Using RNase T1 (navy) and the combination of RNase T1 and RNase 4 coverages (teal). RNase T1 and RNase 4 sequence coverages are the average of two replicates per enzyme. *Data collected in this study was compared to bottom-up IP-RP-MS/MS of *E. coli* total tRNA using two RNase enzymes: RNase T1, cusativin, or MC1 reported previously^20^.

Figure 4 demonstrates the increase in sequence coverage when using folded RNase T1 and complete RNase 4 in orthogonal digestions. For example, tRNA^Glu^ exhibits a small increase in sequence coverage, and tRNA^SerGGA^ also had a large increase in sequence coverage through the addition of RNase 4. For tRNA^Glu^, RNase T1 and RNase 4 had an individual coverage of 94.7% and 100% respectively (**Figure 6A-D**). Thus, the sequence coverage of these two enzymes results in a combined sequence coverage of 100% and result in high sequencing depth across the tRNA body. Other tRNAs had a drastic improvement on sequence coverages, such as tRNA^SerGGA^ where RNase T1 and RNase 4 had 55.7% and 95.5% sequence coverage, respectively. Additionally, some tRNAs did not have an improvement in sequence coverages with RNase 4; however, these tRNAs had an improved sequencing depth, where positions across the tRNA have multiple oligonucleotides mapped, by incorporating an additional enzyme (e.g. Arg^ICG^ and Gln^CUG^) (**Figure 5, SI Table 4, 6, 10**).

The combined sequence coverages we calculated in a total tRNA pool using folded RNase T1 and unfolded (complete) RNase 4 digestions are on average 17.5% higher than previously reported sequence coverages using IP-RP for *E. coli* total tRNA (**Figure 5, SI Table 9**)^20^. This report using IP-RP is, to our knowledge, the only study reporting sequence coverages of *E. coli* total tRNA using multiple RNases and as such is a benchmark for comparison for HILIC-MS/MS and incorporating RNase 4 into the RNA modification mapping workflow. Furthermore, incorporating RNase 4 into this workflow results in higher sequence coverages than those obtained using IP-RP on unfolded (complete) RNase T1 and MC1 (U-specific) or cusativin (C-specific) digestions for 29 of the 35 isoacceptors with an increased sequence coverage ranging from 2.8 - 58% (**Figure 5, SI Table 9**). Therefore, our data suggest that incorporating RNase 4 into total tRNA modification mapping workflows can provide orthogonal data to the currently used enzymes and can increase sequence coverages due to its dinucleotide cleavage sites.

### RNA modification crosstalk can be detected using bottom-up LC-MS/MS

Thus far, bottom-up LC-MS/MS workflows to map RNA modifications in total tRNAs has been limited to wildtype tRNAs and its utility to study enzyme knockouts has yet been explored. RNA modification crosstalk, where the installation of one RNA modification influences the incorporation of another RNA modification, has been well documented^11,30–34^. However, this type of analysis is typically limited to nucleoside LC-MS/MS^11,30,31^. Here, RNA is digested to the nucleoside level, and the global abundance of modified nucleosides is quantified using LC-MS/MS. While this assay can provide insightful quantitative information about how RNA modifying enzymes change the global abundance of RNA modifications, it removes all sequence information. Thus, pinpointing the site of where modification levels are changing can be challenging if the modification of interest is on multiple tRNAs or located at various sites across one or multiple tRNAs. To do this, nucleoside LC-MS/MS must be coupled with an orthogonal method such as nanopore sequencing to see site specific changes in modification status^11^.

To test if RNA modification crosstalk can be detected using bottom-up LC-MS/MS, we digested *E. coli* total tRNA purified from *E. coli* K-12 lacking TrmA, the methyltransferase responsible for the incorporation of 5-methyluridine (m^5^U) at position 54 of tRNAs. m^5^U54 is a conserved RNA modification involved in RNA modification circuits, where the incorporation of m^5^U54 influences the modification of s^4^U8 and acp^3^U47 in tRNA^Phe^. In tRNA^Phe^ purified from *E. coli* lacking TrmA, it has been shown that the nucleoside abundance of s^4^U8 increases, whereas the abundance of acp^3^U47 decreases^30,31^. Our data collected from tRNAs grown in wildtype *E. coli* cells had significant depth on tRNA^Phe^, suggesting that our LC-MS/MS modification mapping method might be useful for directly detecting this RNA modification circuit (**Figure 7**). In tRNA^Phe^ from *trmA*Δ cells, all oligonucleotides with coverage at position 54 map back to uridine rather than m^5^U, validating the *trmA*Δ knockout (**SI Tables 11, 12**). In the m^5^U54 modification circuit in tRNA^Phe^, the abundance of s^4^U increases, which cannot be directly quantified from this assay due to differences in detected oligonucleotides across the two RNase digestions and a lack of an internal standard to normalize the signal to. However, the abundance of modifications that decrease due to modification crosstalk, such as acp^3^U47 could be detected as we would expect to detect modified and unmodified oligonucleotides mapping to position 47. Nucleoside LC-MS/MS has previously revealed that the abundance of acp^3^U47 decreases by 50% in tRNA^Phe^ from *trmA*Δ cells^30,31^. Therefore, we can anticipate a mixture of oligonucleotides with position 47 containing acp^3^U and U in our LC-MS/MS data. We searched both RNase T1 and RNase 4 digestions for oligonucleotides matching either acp^3^U47 and U47 for tRNA^Phe^ and detected oligonucleotides that mapped back to U47 and acp^3^U47 (**Figure 7, SI Table 11, 12**). Despite the oligonucleotide that maps U47 being a shorter oligonucleotide, we are confident that this arises from the modification circuitry of m^5^U as the oligonucleotide mapping U47 is only identified in RNase digest from tRNAs purified from cells lacking trmA. While the modified oligonucleotide is detected in digested tRNAs purified from WT cells, these two oligonucleotides retain differently on the HILIC column. This reveals that the U47 oligonucleotide is a result of m^5^U-acp^3^U crosstalk, rather than an artifact due to in-source fragmentation which can result in modifications prematurely fragmenting off the oligonucleotide before MS detection (**SI Table 3, 11**). Furthermore, when compared to a theoretical MS/MS spectra, the oligonucleotide with U47 has a shift in mass of −101 Da in C-type ions, characteristic of the presence of U47 (Figure 7C). Together, the differences in HILIC retention between acp^3^U47 and U47 oligonucleotides and the shift in mass in the MS/MS spectra make us confident that this oligonucleotide is a result of modification crosstalk due to the loss of m^5^U by knocking out trmA.

**Figure 6.**
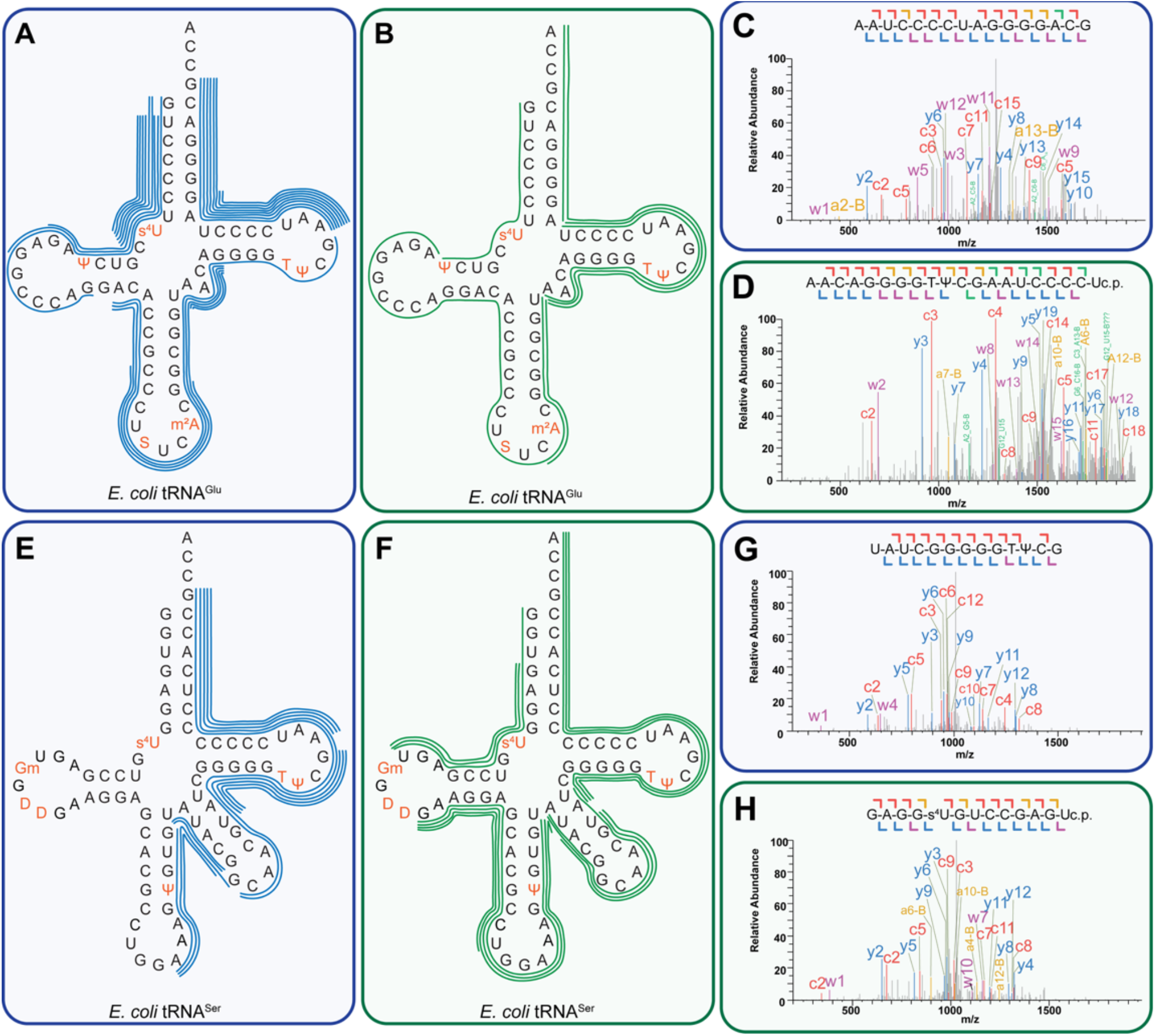
Combined RNase T1 and RNase 4 sequence coverages. A) RNase T1 sequence coverage of tRNA^Glu^. Each line corresponds to a detected oligonucleotide. B) RNase 4 sequence coverage of tRNA^Glu^. C) MS/MS spectrum of tRNA^Glu^ oligonucleotide generated from RNase T1 digestion that maps from the T-loop to the acceptor stem (A58-G73). This oligonucleotide has a m/z of 1278.684 and has a terminal 3’ OH end. D) MS/MS spectrum of tRNA^Glu^ oligonucleotide that maps across the variable loop and T-loop (A46-U65). This oligonucleotide has a m/z of 1609.212 and a terminal 3’ cyclic phosphate. E) RNase T1 sequence coverage of tRNA^Ser(GGA)^. Each line corresponds to a detected oligonucleotide. F) RNase 4 sequence coverage of tRNA^Ser(GGA)^. G) MS/MS spectrum of tRNA^Ser(GGA)^ oligonucleotide generated from RNase T1 digestion that maps from the variable loop to the T-loop (U57-G69). This oligonucleotide has a m/z of 1045.640 and has a terminal 3’ OH end. H) MS/MS spectrum of tRNA^Ser(GGA)^ oligonucleotide from RNase 4 digestion that maps from the acceptor stem to the D-loop (G4-U16). This oligonucleotide has a m/z of 1067.375 and has a terminal 3’ cyclic phosphate end.

**Figure 7.**
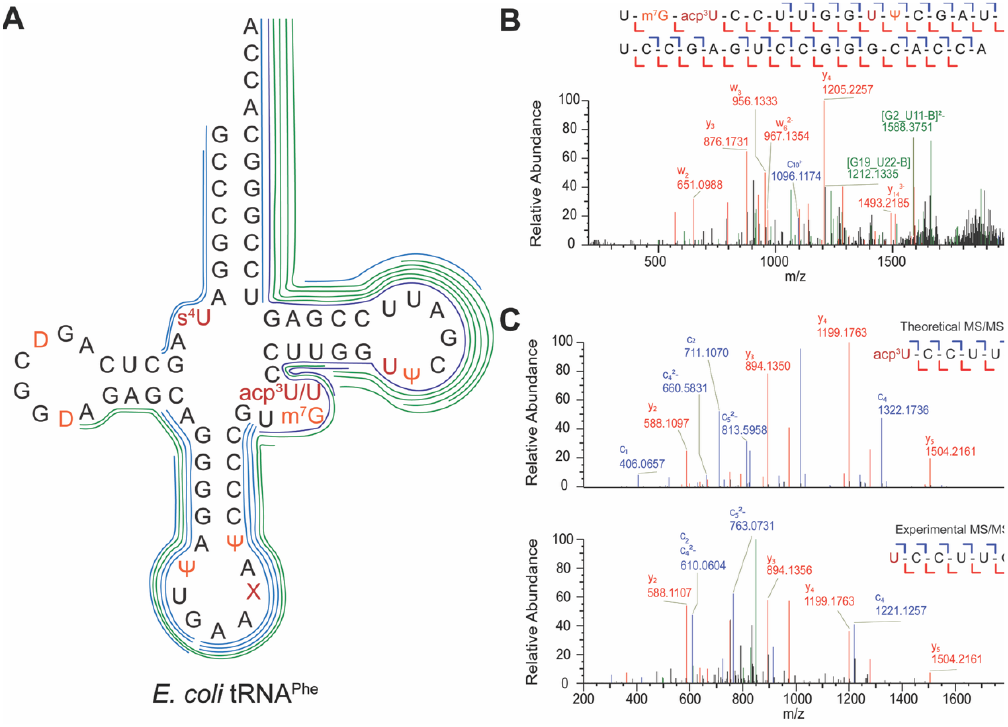
Detected modification crosstalk in E. coli tRNA^Phe^ between m^5^U54 and acp^3^U47. A) RNase T1 and RNase 4 detected tRNA^Phe^ oligonucleotides from trmAΔ cells responsible for m^5^U54. Modification labeled X represents ms^2^i^6^A. RNase T1 oligonucleotides are represented in blue. RNase 4 oliognucleotides are represented in green. The dark blue oligonucleotides MS/MS are shown in panels B and C. B) MS/MS fragmentation of oligonucleotide mapping acp^3^U47 and U54 with a m/z 1712.895 from RNase T1 digest. D) MS/MS fragmentation of oligonucleotide mapping U47 with a m/z 904.620 from RNase T1 digest.

## DISCUSSION

Here we demonstrate the application of RNase 4 in bottom-up HILIC-MS/MS RNA modification mapping of *E. coli* total tRNA. Despite the highly structured and modified nature of tRNAs, RNase 4 produced oligonucleotide digestion products in ranges compatible with HILIC-MS/MS. The majority of the detected digestion products were 10-20 nt long and can be resolved well using HILIC, easily ionized by ESI, and uniquely map back to a specific tRNA. Additionally, we have shown that the folded digestion scheme we developed for commercial *S. cerevisiae* tRNA can be applied to tRNA of other organisms with the optimization of enzyme concentrations. Furthermore, this suggests that the folded digestion scheme can be used to map modifications with high sequence coverages in any RNA that is highly structured (i.e. rRNA).

Using HILIC-MS/MS, our results are comparable with data collected previously using IP-RP for *E. coli* total tRNA. Despite the excellent resolution IP-RP can offer for oligonucleotide separations, our results have high sequence coverage and depth across the *E. coli* isoacceptors, in similar analysis times. While our method uses a larger RNA input (10 μg vs 225 μg), HILIC negates the need for dedicated instruments necessary for IP-RP analysis and provides an alternative mode of chromatography for the analysis of oligonucleotides. Our approach showed sufficient depth to identify previously unrecognized modification sites, including tRNA^Gly(mnm5UCC)^ (**Figure 4 C,D**), even in the well-characterized model organism *E. coli*, whose tRNA modification profiles have long been considered “established.”

Furthermore, our work to improve HILIC-MS/MS RNA modification mapping will be beneficial for interpreting orthogonal sequencing methods that do not directly report on modification identity, including nanopore direct RNA sequencing (DRS) data which required substantially less RNA to perform (300 ng). Here, we combined nanopore DRS analysis of Δ*thiI* with LC-MS/MS sequencing to confidently assign a predicted s^4^U modification a position 8 in Arg^UCU^ tRNAs^48^. In addition to improving coverage across many tRNAs, we have also been the first to show the application of bottom-up LC-MS/MS to detect RNA modification crosstalk. RNA modification crosstalk has mainly been studied by using nucleoside LC-MS/MS which eliminates all sequence context, or by direct RNA sequencing which needs an orthogonal method for validation. Therefore, these results suggest that bottom-up LC-MS/MS can detect RNA modification circuits/crosstalk and should be implemented with direct RNA sequencing to provide orthogonal validation.

## ACKNOWLEDGEMENTS

We thank Roja Gundepudi (University of Michigan) for the growth and purification of total tRNA used in the manuscript. We thank members of the Koutmou and Kennedy Labs for their comments on the manuscript. Features of the graphical abstract were made with BioRender (agreement # OZ28XCRZQ0)^42^.

## AUTHOR CONTRIBUTIONS

Kaley M Simcox: Conceptualization, Formal analysis, Investigation, Methodology, Validation, Writing—original draft, Writing—review & editing.

Maddy Zamecnik: Formal analysis, Investigation, Methodology, Validation, Visualization, Writing—original draft, Writing—review & editing.

Robert T. Kennedy: Conceptualization, Formal analysis, Funding acquisition, Methodology, Supervision, Validation, Writing—review & editing.

Kristin S. Koutmou: Conceptualization, Formal analysis, Funding acquisition, Methodology, Supervision, Validation, Writing—original draft, Writing—review & editing.

## SUPPLEMENTARY DATA

Supplementary Data are available at XXX online.

## DATA AVAILABILITY

The RNA mass spectrometry data have been deposited to the ProteomeXchange Consortium via the PRIDE [1] partner repository with the dataset identifier PXD068701.

