## Supplementary Figures for "RNase 4 improves bottom-up modification mapping of *E. coli* total tRNAs using HILIC-MS/MS"

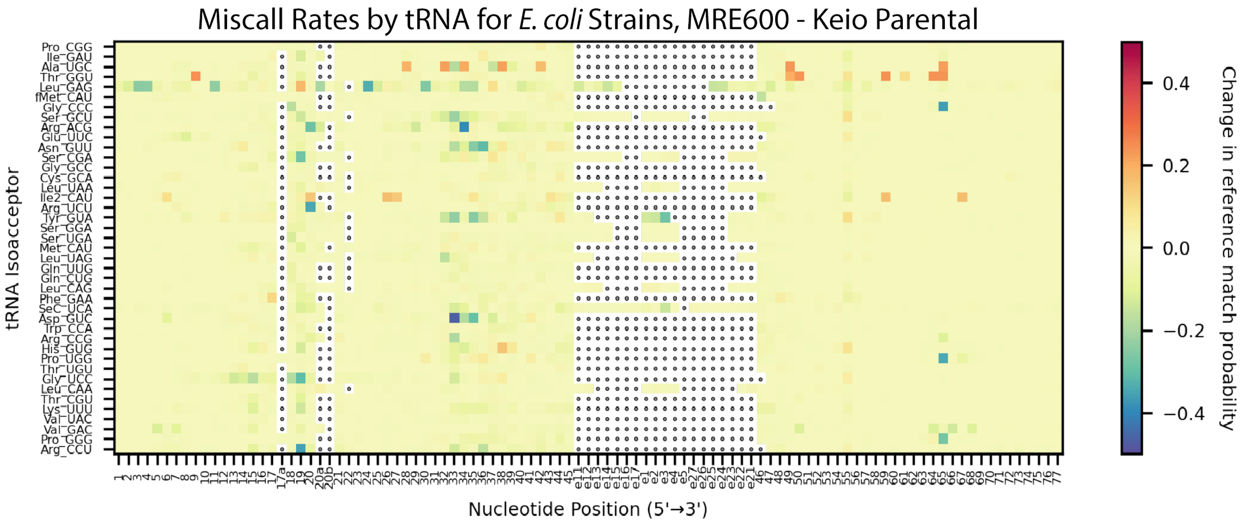


**SI Figure 1:** Heatmap showing the change in modification status between total tRNA samples from two different strains of *E. coli*, the MRE600 and K-12 strains, sequenced using Oxford Nanopore direct RNA sequencing. Using miscalls as a proxy for modification status, red indicates positions with higher stoichiometry modifications in the MRE600 strain, and blue represents higher modification stoichiometry in the K-12 parental strain.

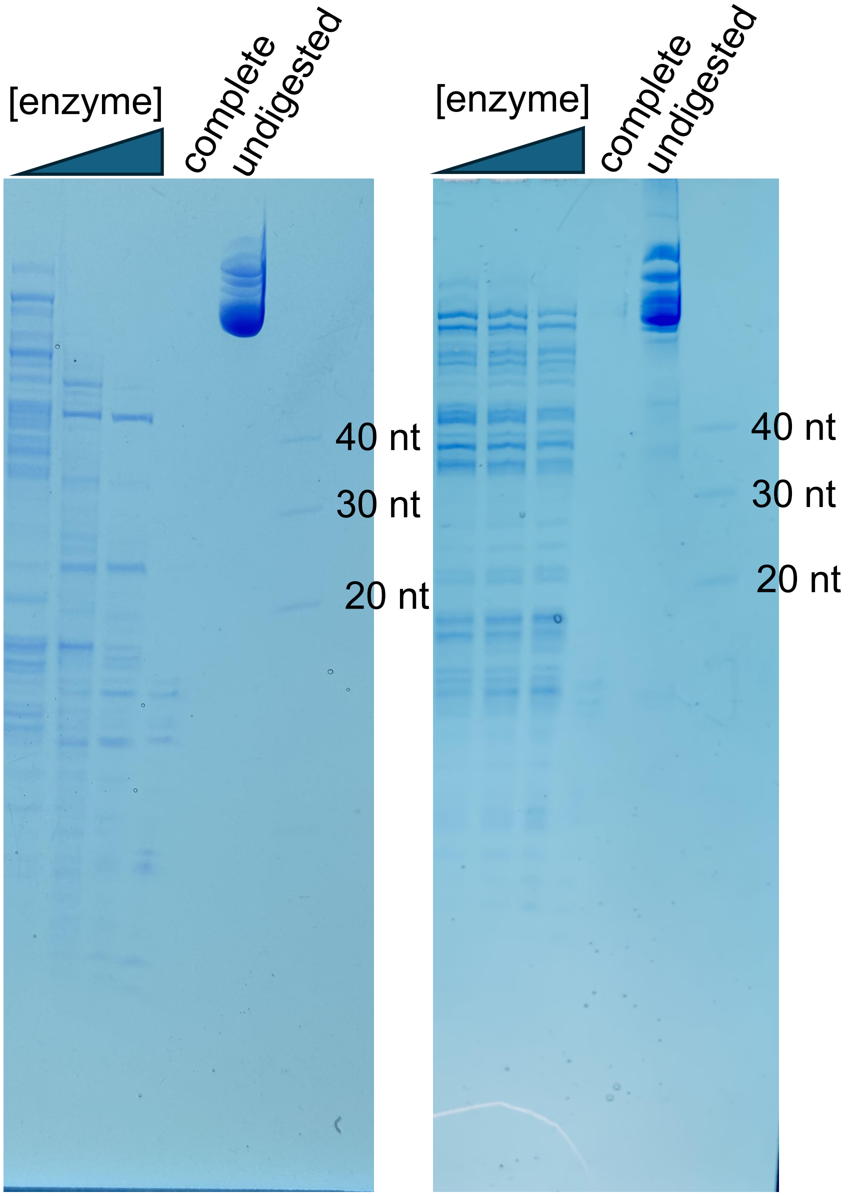


**SI Figure 2:** Urea-PAGE with folded RNase T1 digestions using commercially available *E. coli* total tRNA (Roche #10109541001) and *E. coli* total tRNA purified in our lab. **Left -** Purified *E. coli* total tRNA. Lane 1: RNA ladder, 40 nt, 30 nt, 20 nt, 10 nt. Lane 2: undigested tRNA. Lane 3: complete or unfolded RNase T1 digestion. Lane 4 and 7: 25 U/ug RNase T1. Lane 5 and 8: 12 U/ug RNase T1.Lane 6 and 9: 6 U/ug RNase T1. **B.** Commercially available *E. coli* total tRNA. Lane 1: RNA ladder, 40 nt, 30 nt, 20 nt, 10 nt. Lane 2: undigested tRNA. Lane 3: unfolded or complete RNase T1 digestion. Lane 4 and 7: 25 U/ug RNase T1. Lane 5 and 8: 12 U/ug RNase T1. Lane 6 and 9: 6 U/ug RNase T1.

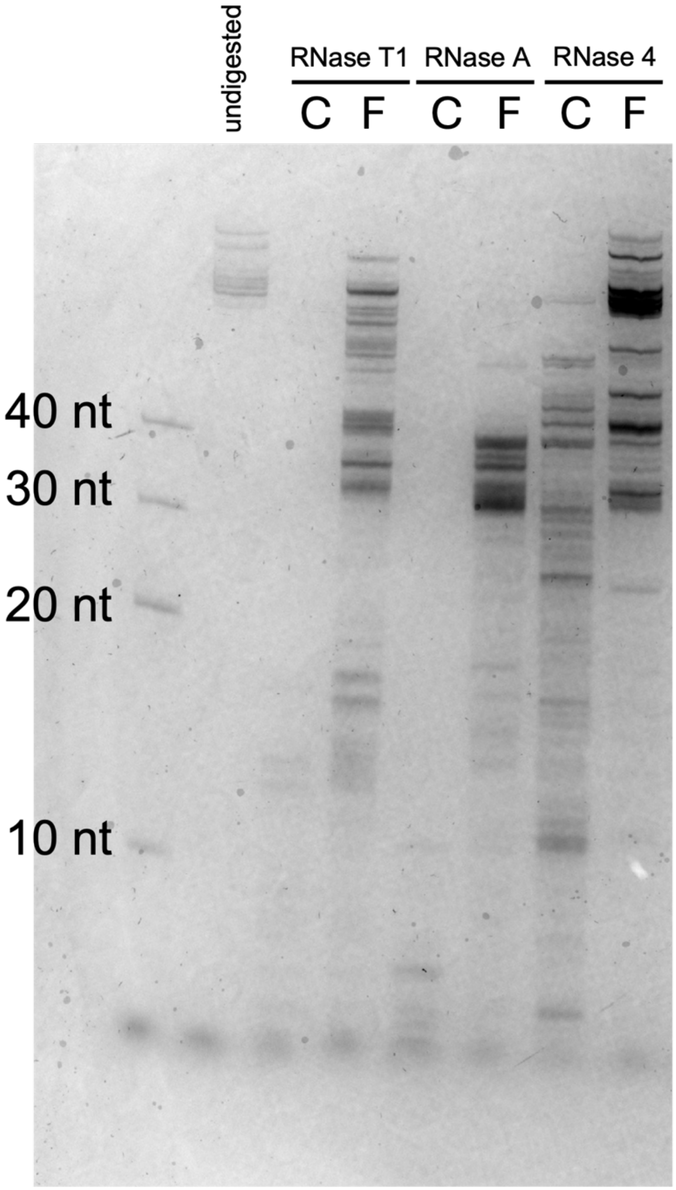


**SI Figure 3:** Urea-PAGE with folded vs unfolded digestions stained with SYBR Gold (Invitrogen). Lane 1: RNA ladder, 40 nt, 30 nt, 20 nt, 10 nt. Lane 2: undigested RNA. Lane 3: complete RNase T1 digestion. Lane 4: Folded RNase T1 digestion. Lane 5: complete RNase A digestion. Lane 6: Folded RNase A digestion. Lane 7: complete RNase 4 digestion. Lane 8: folded RNase 4 digestion.

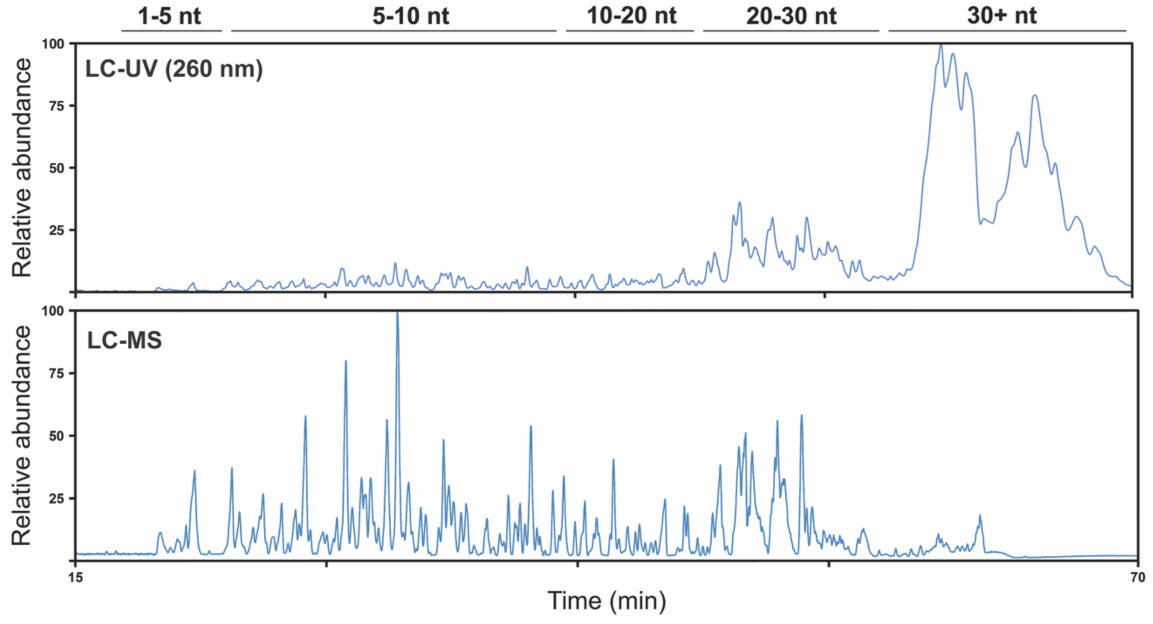


SI Figure 4: Comparison of LC-UV and LC-MS signal for *E. coli* total tRNA digestion. UV data is collected prior to MS detection. Top panel – LC-UV trace of *E. coli* tRNA RNase T1 digestion monitored at 260 nm. Bottom panel – Base peak chromatogram of *E. coli* tRNA RNase T1 digest.

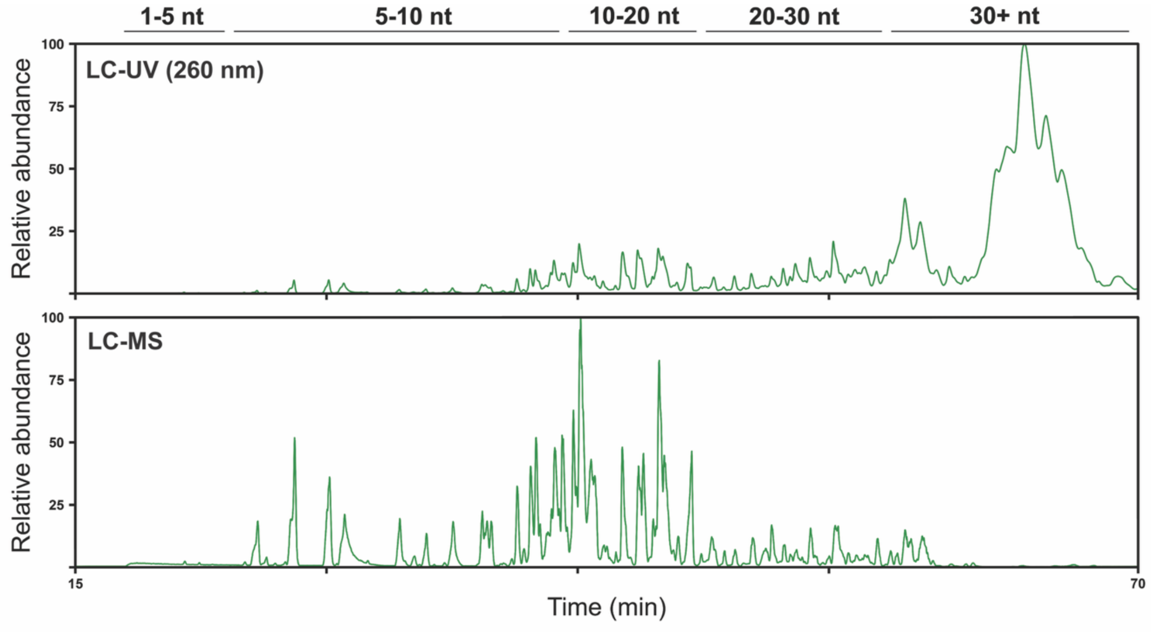


SI Figure 5: Comparison of LC-UV and LC-MS signal for *E. coli* total tRNA RNase 4 digestion. UV data is collected prior to MS detection. Top panel – LC-UV trace of *E. coli* tRNA RNase 4 digestion monitored at 260 nm. Bottom panel – Base peak chromatogram of *E. coli* tRNA RNase 4 digest.

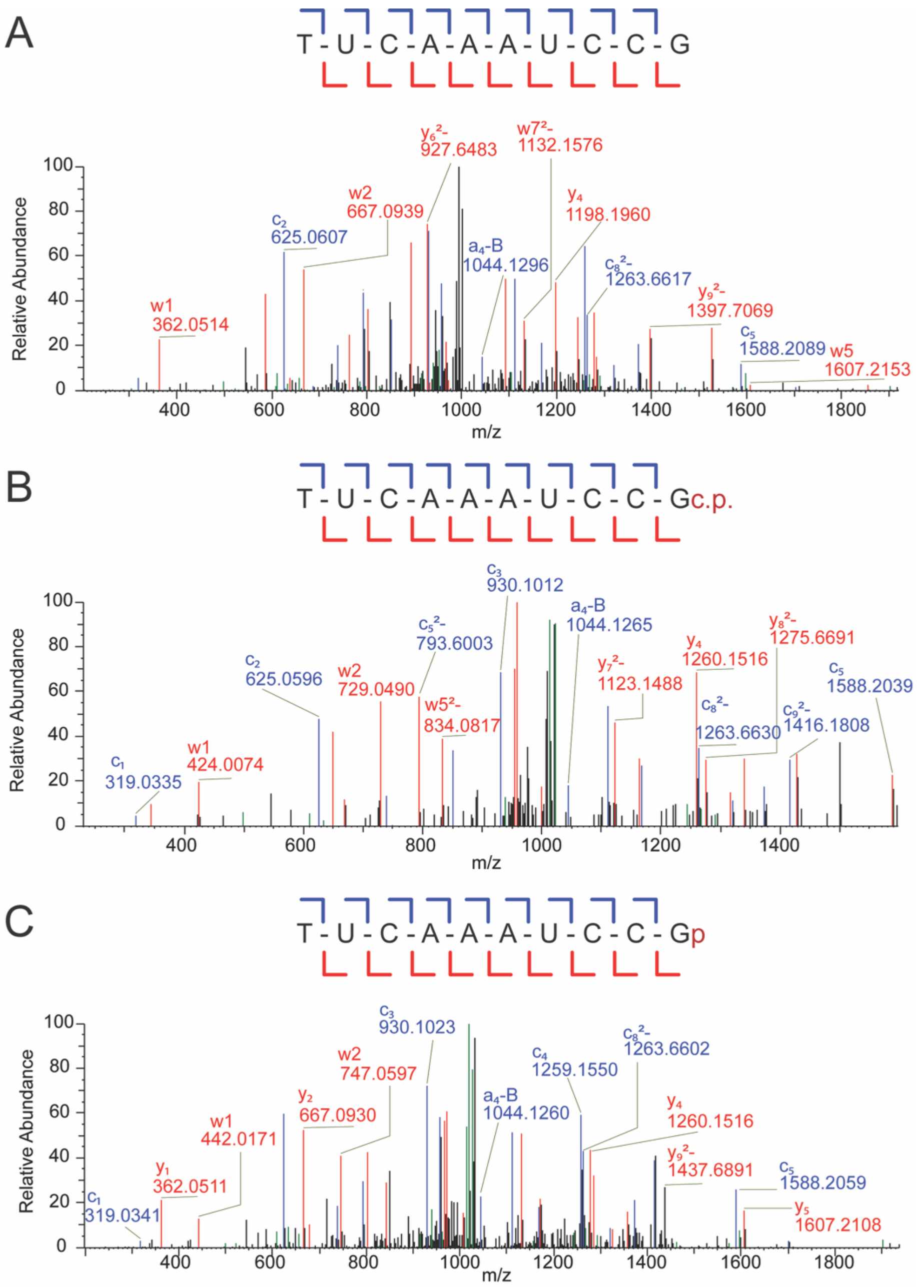


**SI Figure 6: Variable 3′ phosphate ends, from T1 digest tRNA Ini**

B-D) MS/MS spectra of oligonucleotides mapping from position XX to postion XX in the T-loop of tRNA$^{Ini}$ (shown in green) with varying phosphate ends. The colors in the MS/MS spectra correspond to the type of ion generated through ion-trap CID. The colors are as follows C-type (red), Y-type (blue), a-B-type (yellow), W-type purple. B) MS/MS spectrum with a terminal 3′ OH. C) MS/MS spectrum with a terminal 3′ cyclic phosphate. D) MS/MS spectrum with a terminal 3′ linear phosphate.

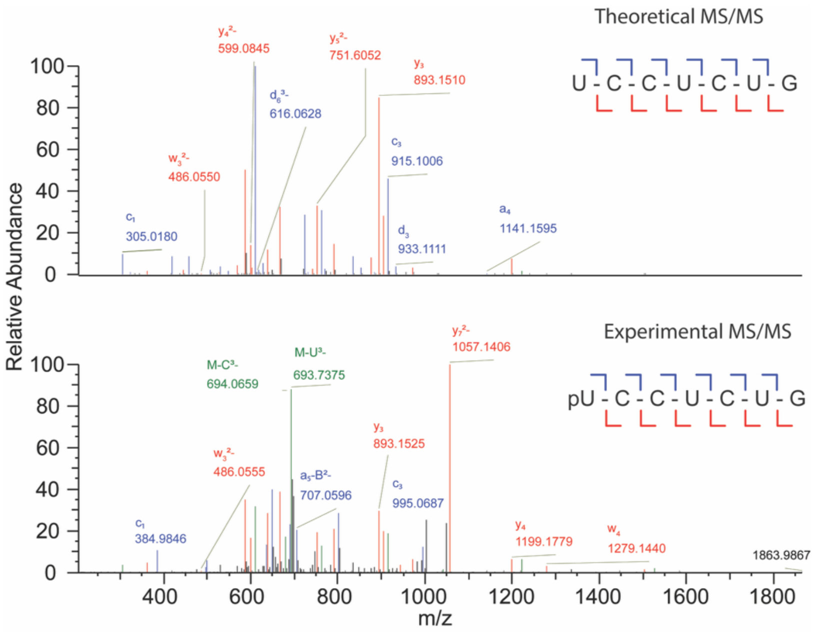


**SI Figure 7: 5′ phosphates – Asn_QUU T1 digest**

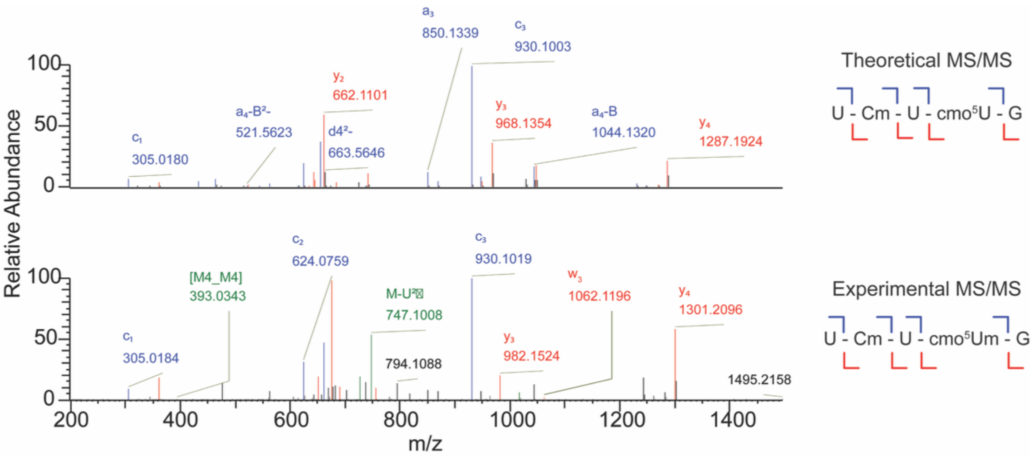


**Figure 8: cmo5Um Ser_UGA T1**

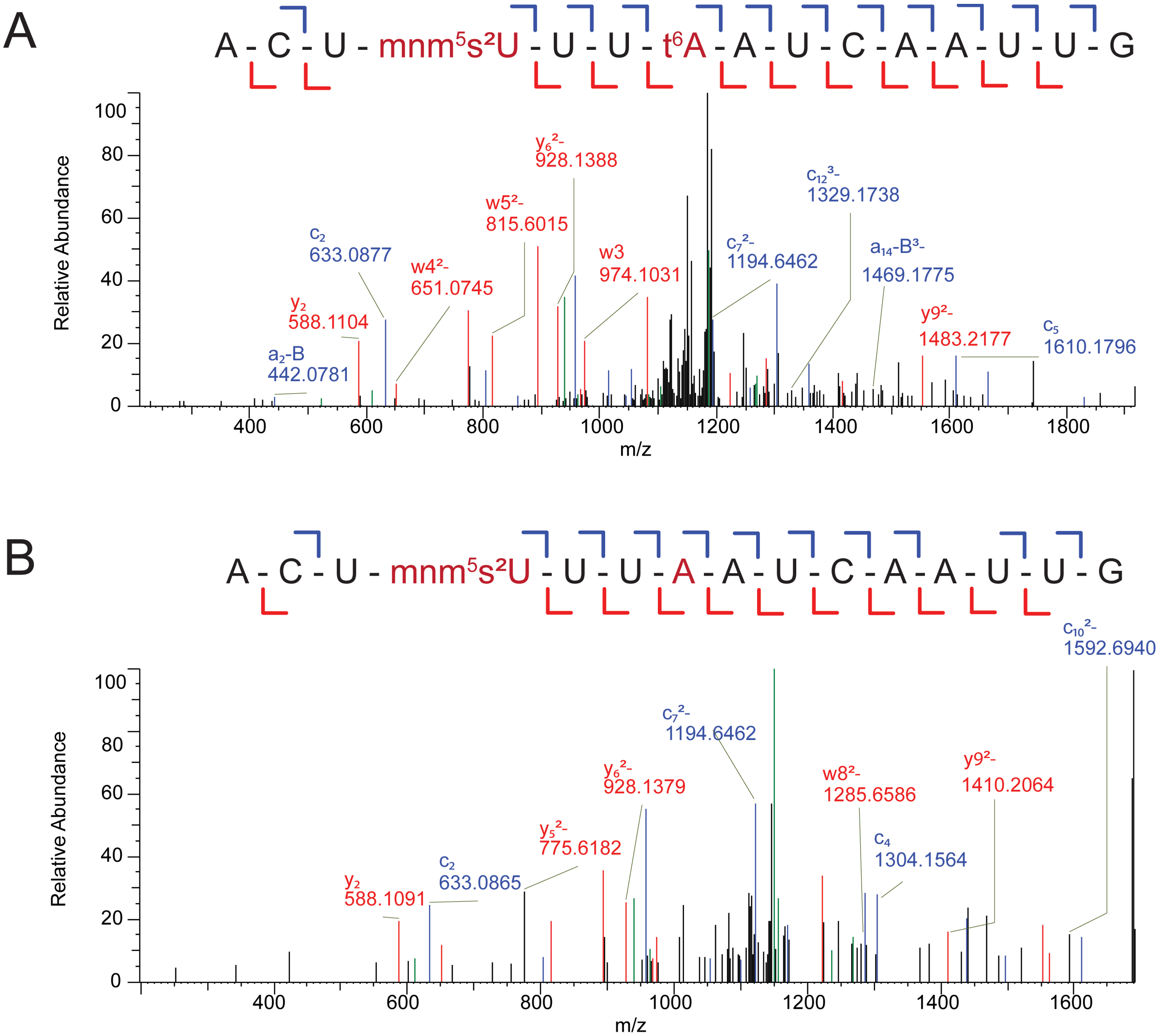


**SI Figure 9: ASL mods all from T1 digest Lys_SUU**

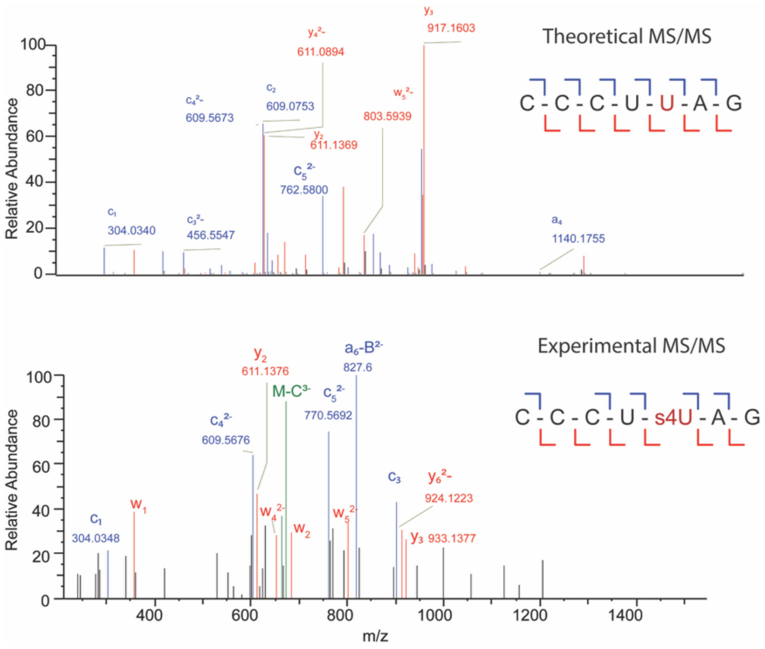


**SI Figure 10: s4U8 Arg UCU T1**

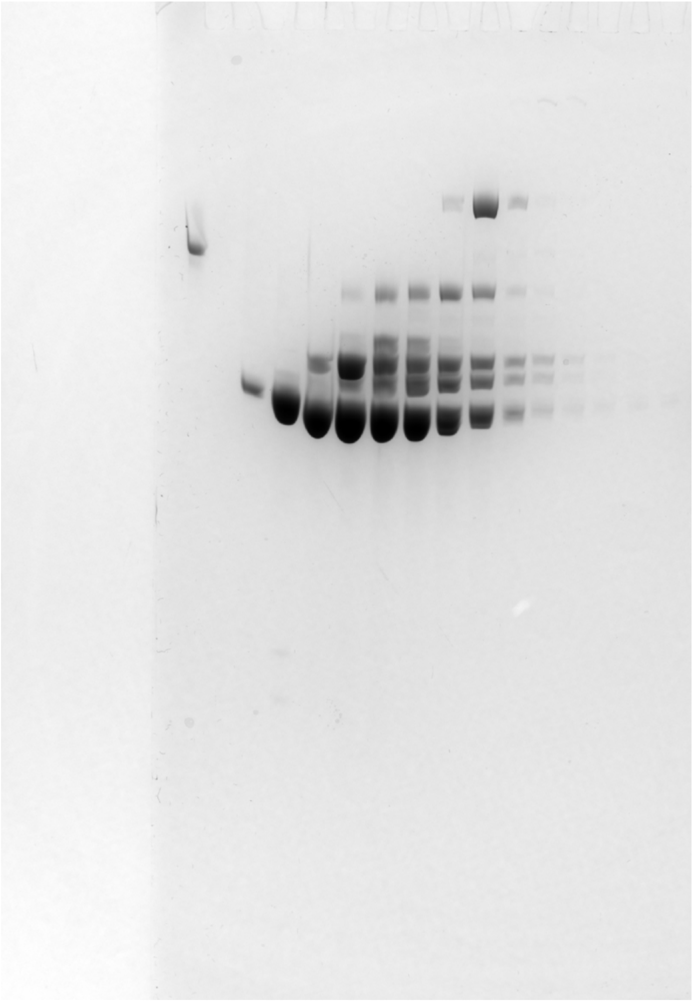

**SI Figure 11: FPLC fractions from total RNA purified using strong anion exchange ran on urea-PAGE and stained with SYBR Gold.** Lane 1: 5S rRNA maker (120 nt). Lanes 2-12: FPLC fractions from strong anion exchange on a 6-mL Cytiva Resource Q column. The first 3 FPLC fractions were pooled to have total tRNA.
